# Belowground changes to community structure alter methane-cycling dynamics in Amazonia

**DOI:** 10.1101/2020.03.10.984807

**Authors:** Kyle M. Meyer, Andrew H. Morris, Kevin Webster, Ann M. Klein, Marie E. Kroeger, Laura K. Meredith, Andreas Brændholt, Fernanda Nakamura, Andressa Venturini, Leandro Fonseca de Souza, Katherine L. Shek, Rachel Danielson, Joost van Haren, Plinio Barbosa de Camargo, Siu Mui Tsai, Fernando Dini-Andreote, José M. S. de Mauro, Klaus Nüsslein, Scott Saleska, Jorge L. M. Rodrigues, Brendan J. M. Bohannan

## Abstract

Amazonian rainforest is undergoing increasing rates of deforestation, driven primarily by cattle pasture expansion. Forest-to-pasture conversion has been associated with changes to ecosystem processes, including substantial increases in soil methane (CH_4_) emission. The drivers of this change in CH_4_ flux are not well understood. To address this knowledge gap, we measured soil CH_4_ flux, environmental conditions, and belowground microbial community attributes across a land use change gradient (old growth primary forest, cattle pasture, and secondary forest regrowth) in two Amazon Basin regions. Primary forest soils exhibited CH_4_ uptake at modest rates, while pasture soils exhibited CH_4_ emission at high but variable rates. Secondary forest soils exhibited low rates of CH_4_ uptake, suggesting that forest regrowth following pasture abandonment could reverse the CH_4_ sink-to-source trend. While few environmental variables were significantly associated with CH_4_ flux, we identified numerous microbial community attributes in the surface soil that explained substantial variation in CH_4_ flux with land use change. Among the strongest predictors were the relative abundance and diversity of methanogens, which both increased in pasture relative to forests. We further identified individual taxa that were associated with CH_4_ fluxes and which collectively explained ~50% of flux variance. These taxa included methanogens and methanotrophs, as well as taxa that may indirectly influence CH_4_ flux through acetate production, iron reduction, and nitrogen transformations. Each land type had a unique subset of taxa associated with CH_4_ fluxes, suggesting that land use change alters CH_4_ cycling through shifts in microbial community composition. Taken together, our results suggest that changes in CH_4_ flux from agricultural conversion could be driven by microbial responses to land use change in the surface soil, with both direct and indirect effects on CH_4_ cycling. This demonstrates the central role of microorganisms in mediating ecosystem responses to land use change in the Amazon Basin.

## INTRODUCTION

After a decade of slowing rates of deforestation, the Amazon rainforest is again undergoing high rates of deforestation, driven primarily by agricultural expansion for cattle pasture (Laurance *et al*. 2014; Barlow *et al*. 2019). Such forms of environmental change are known to alter belowground microbial biodiversity (Rodrigues *et al*. 2013; Mueller *et al*. 2016; Meyer *et al*. 2017) as well as microbially-mediated biogeochemical cycles (Neill *et al*. 1997b, 2005; Verchot *et al*. 1999), including the methane (CH_4_) cycle. Rainforest soils in the western Amazon Basin switch from acting as a sink for atmospheric CH_4_ to a persistent source of CH_4_ following conversion (Steudler *et al*. 1996; Fernandes *et al*. 2002), and little is known about whether the CH_4_ sink capacity returns following pasture abandonment and secondary forest regeneration. This sink-to-source phenomenon has also been documented in the Eastern Amazon (Keller *et al*. 1986; Verchot *et al*. 2000), suggesting a general functional response to cattle pasture establishment. This is of concern considering recent increases in agricultural conversion throughout the Amazon Basin (Carvalho *et al*. 2019), and the fact that CH_4_ is a potent greenhouse gas, with roughly 34 times the global warming potential of CO_2_ over a 100-year timeframe (Myhre *et al*. 2013). Although responses of belowground microbial communities and CH_4_ flux to land use change have both been documented in the Amazon Basin (Keller *et al*. 1986; Steudler *et al*. 1996; Verchot *et al*. 2000; Fernandes *et al*. 2002; Meyer *et al*. 2017), the relationship between these two responses is not well understood, in part because no study has measured microbial community attributes and CH_4_ flux simultaneously.

Soil CH_4_ flux results from two counter-acting microbial processes: CH_4_ production (methanogenesis) and CH_4_ consumption (methanotrophy) (Conrad 2009). Methanogens are Archaea that anaerobically produce CH_4_ using either acetate, methylated compounds, formate, or H_2_ and CO_2_ (Hedderich & Whitman 2013). Methanogens have been shown to increase in relative abundance following conversion to cattle pasture, as well as undergo compositional changes that may indicate a shift in the predominance of methanogenic pathways (Meyer *et al*. 2017). Aerobic methanotrophs are Bacteria in the Alpha- and Gamma-Proteobacteria and Verrucomicrobia that consume CH_4_ via the serine, ribulose monophosphate (RuMP), or Calvin-Benson-Bassham pathways, respectively (Knief 2015). Methanotrophs have also been reported to strongly respond to land use change in the Amazon, including decreases in population abundance and alterations to community composition (Meyer *et al*. 2017).

Methanogens and methanotrophs are the only groups to directly cycle CH_4_, but these organisms form complex ecological interactions with other community members and this may influence the rate or directionality of CH_4_ flux. For example, methanogens depend on metabolic byproducts (e.g. H_2_ and CO_2_, or acetate) derived from the activity of other community members such as acetogens or fermentative bacteria (Müller & Frerichs 2013). Methanogens are often outcompeted by other community members for these substrates when more thermodynamically favorable terminal electron acceptors are available, including NO_3^-^_, NO_2_, and Fe (II), (Cord-Ruwisch *et al*. 1988; Chen & Lin 1993; Klüber & Conrad 1998). The activity of aerobic methanotrophs can also depend on community interactions, such as competition for O_2_ or soil nitrogen (N) (Bodelier *et al*. 2000; Bodelier & Steenbergh 2014; Ho *et al*. 2016), or predation by protozoa or viruses (Tyutikov *et al*. 1980; Murase & Frenzel 2008). To date, few have sought to relate broader community interactions to soil CH_4_ emissions.

There is growing interest in better understanding ecosystem functions using microbial community measurements (McGuire & Treseder 2010; Bier *et al*. 2015; Hall *et al*. 2018), but attempts have generated mixed results (Rocca *et al*. 2015; Graham *et al*. 2016; Louca *et al*. 2018). Microbial taxa can be artefactually related to CH_4_ flux due to covariation with environmental conditions that alter function, or through spatial auto-correlation (Legendre 1993), and this covariance structure could blur the connection between communities and function (Morris *et al*. 2019). Accounting for covariance structure has been shown to aid in detecting microbial taxa or community attributes that are causally associated with CH_4_ processes (Meyer *et al*. 2019). One way to do so uses principle components analysis to derive environmental, spatial, and community structure covariates for incorporation into statistical models (Price *et al*. 2006; Morris *et al*. 2019). Applying this approach could help clarify the relationship between environmental change and community functional responses, especially in ecosystems such as those of the Amazon Basin, where many variables exhibit change following ecosystem conversion.

This study focuses on a gradient of land use change in two regions of the Amazon Basin. We combine measurements of *in situ* CH_4_ flux, soil chemistry, and microbial community structure across primary rainforest, cattle pasture, and secondary forest (derived from abandoned cattle pasture). We first ask how land use change alters soil CH_4_ flux and the community structure of bacteria and archaea (including CH_4_-cycling organisms). We then investigate the relationships between environmental variables, microbial community attributes, and CH_4_ flux, in order to identify mechanisms that link land use change to changes in CH_4_ flux. Our study provides an important window into a poorly understood phenomenon that is likely to become increasingly common throughout the Amazon Basin if rates of land use change continue to increase.

## METHODS

### Site description, sampling design, sampling dates

Our study was performed in two regions of the Amazon Basin: the state of Rondônia in the Western Amazon, and the state of Pará in the Eastern Amazon. Both states have experienced the highest rates of forest loss in Brazil, largely driven by agricultural expansion for cattle ranching (Soares-Filho *et al*. 2006; Ometto *et al*. 2011; Carvalho *et al*. 2019). In Rondônia, we surveyed three primary forest sites, three cattle pasture sites, and two secondary forest sites, totaling 39 sampling locations, all in or directly adjacent to Fazenda Nova Vida, about 250 km south of Porto Velho. The climate at Fazenda Nova Vida is humid tropical, and receives 2200 mm annual mean precipitation (Steudler *et al*. 1996; Alvares *et al*. 2013). Soils are red-yellow podzolic latosol with sandy clay loam texture, and are described in detail elsewhere (Neill *et al*. 1997a). Vegetation type is open moist tropical forest with palms, and is described elsewhere (Pires & Prance 1985). In Pará we surveyed two primary forest sites, three cattle pasture sites, and three secondary forest sites, totaling 33 sampling locations. Pará sites were in or around Tapajós National Forest, which receives roughly 2000 mm annual mean precipitation. Soils there have been characterized as ultisols and oxisols in flat areas, and inceptisols in areas with topographic relief, and have been further described alongside floristic descriptions elsewhere (Parrotta *et al*. 1995; Silver *et al*. 2000; Espírito-Santo *et al*. 2005; Keller *et al*. 2005). We strove to sample forests, pastures, and secondary forests equally, but faced restrictions due to varying land ownership and logistical issues. At each site, we established a 200 m transect and performed paired sampling of gases and soil at 50 m intervals, with 5 locations for measurements and sampling per transect. Sampling in Pará and Rondônia took place during wet season periods, in June 2016 and March/April of 2017, respectively. GPS coordinates of each sampling point can be found in Supplementary Table 1.

At each sampling location soil CH_4_ flux was measured in real time using a field-deployable Fourier-transform infrared spectrometer (Gasmet, DX 4015, Vantaa, Finland) connected to a flow-through soil flux chamber in a closed recirculating loop. Soil collars (aluminum, inner area of 284 cm^2^) were installed roughly 5 cm into the soil surface at least 20 minutes before CH_4_ concentration measurements began. Soil flux chambers were connected via inlet and outlet ports to the CH_4_ analyzer and were placed on the soil collars. CH_4_ fluxes were determined by the rate of accumulation or removal of CH_4_ in the flux chamber headspace over a 30-minute period. Trends in CH_4_ concentration over time varied from linear to non-linear. If trends in CH_4_ concentration over time were linear, a linear model was used to calculate flux. If trends were non-linear, we used the linear portion of the data near the time of chamber placement to calculate flux (Salimon *et al*. 2004; Pirk *et al*. 2016).

Directly following gas flux measurement, soil samples were taken with a sterilized corer (5 cm diameter x 10 cm length) positioned under the chamber and another four cores forming a square around the chamber at ~25 cm distance, to capture community heterogeneity surrounding the chamber area. The five soil cores were emptied into a 4l plastic bag, then mixed by hand from the outside of the bag following root removal. Two 200 g samples of this soil mixture were placed into new sample bags and either frozen for DNA extraction or stored at 4° C for soil chemical analysis.

### Soil chemical analysis

We assessed 19 soil chemical attributes for use as environmental covariates. Soil chemical analyses were performed at the Laboratory of Soil Analysis at “Luiz de Queiroz” College of Agriculture (ESALQ/USP; Piracicaba, Brazil), following the methodology described by (van Raij *et al*. 2001). Soil chemical parameters included pH, organic matter, P, S, K, Ca, Mg, Al, H^+^, Al, sum of bases, cation exchange capacity, base saturation (% V), Al saturation, Cu, Fe, Mn, Zn, and total N. All soil chemical data can be accessed in Supplementary Table 1. For one forest site in Pará, soil chemical data are missing due to a sample transport error. These samples were excluded from microbial analyses requiring environmental covariates, but were included for analyses independent of environmental data (i.e. community structure).

### Soil DNA extraction

Total DNA from each sample was extracted from 0.25 g soil using the DNeasy PowerSoil kit (Qiagen Inc., Valencia, CA, USA) following manufacturer’s instructions. Soil samples from Pará sites required two subtle modifications, based on Venturini *et al*. (2019): 1) vortexing was performed for 15 minutes, instead of 10 minutes, and 2) all incubations steps were at −20° C degrees, instead of 4° C. It is possible that these subtle modifications could influence our results, but they were necessary to obtain quantifiable amounts of DNA, likely due to soil inhibitors such as humic acids. DNA yield from each extraction was fluorometrically quantified (Qubit, Life Technologies, USA).

### Soil prokaryotic community structure assessment

In order to assess the community structure and diversity of soil prokaryotes in each sample, we performed Illumina Miseq 300 basepair paired-end sequencing of the V4 region of the 16S rRNA gene using the 515F - 806R primer combination (Caporaso *et al*. 2011) at the University of Oregon Genomics Core Facility. PCR mixtures were: 12.5 μl NEBNext Q5 Hot Start HiFi PCR master mix, 11.5 μl primer mixture (1.09 μM concentration), and 1 μl of DNA template (total of 17.5 ng DNA per reaction). Reaction conditions were: 98° C for 30s (initialization), 98° C for 10s (denaturation), 61° C for 20s (annealing), and 72° C for 20s (final extension). Reactions were run for 20 cycles and amplicons were purified using 20 μl Mag-Bind RxnPure Plus isolation beads (Omega Bio-Tek, USA). Sequencing libraries were prepared using a dual-indexing approach (Kozich *et al*. 2013; Fadrosh *et al*. 2014), and samples were pooled at equimolar concentration. The final library was sequenced at a concentration of 3.312 ng/μl.

Paired sequence reads were merged using PEAR (version 0.9.10) with default parameters (Zhang *et al*. 2014). Merged reads were filtered by length (retaining read lengths of 230-350 basepairs) and quality (retaining only reads with quality score >30) using Prinseq (Schmieder & Edwards 2011). Filtered sequences were checked for chimeras, denoised, and collected into amplicon sequence variants (ASVs) using DADA2 (Version 1.6) (Callahan *et al*. 2016) implemented in QIIME2 (Bolyen *et al*. 2019). Taxonomy was assigned to ASVs using the RDP naïve Bayesian rRNA classifier Version 2.11 (Wang *et al*. 2007; Cole *et al*. 2014) with training set 16.

### Quantitative PCR of methanogens and methanotrophs

We estimated the abundance of methanogens and methanotrophs using quantitative PCR (qPCR) of marker genes. For methanogens, we targeted the *mcrA* gene using the mlas-mcraRev primer combination (Steinberg & Regan 2008). For methanotrophs, we targeted the *pmoA* gene using the A189 – mb661 primer combination (Bourne *et al*. 2001). DNA from each soil sample was amplified in triplicate using a blocked design whereby all 72 samples (as well as positive and negative controls) were run in a single 96-well plate, repeated three times. Reactions were run on a Bio-Rad CFX96 real-time qPCR instrument (Bio-Rad, USA), using Sso Advanced Universal SYBR Green Supermix reagents (Bio-Rad, USA). Reaction conditions were optimized using an annealing temperature gradient. For each reaction, 2 ng of DNA were used and reactions took place under the following conditions: 98° C 10 minutes (initialization), 98° C 15 seconds (denaturation), 55.6° C 15 seconds (annealing), 72° C 60 seconds (final extension). For both genes, sample amplification was compared to a standard positive control to calculate gene copy number. For *pmoA* the positive control was genomic DNA from *Methylococcus capsulatus* Foster and Davis (ATCC 33009D-5). For *mcrA* the positive control was a *mcrA* copy ligated into a vector. We used LinRegPCR (Ramakers *et al*. 2003; Ruijter *et al*. 2009) to process amplification data, which calculates individual PCR efficiencies. Individual PCR efficiencies were significantly different among regions (Rondônia versus Pará), so the average PCR efficiency for each region was used to calculate gene copy number. To account for plate-to-plate variation (among technical replicates) gene count values for each sample were residualized (by subtracting the mean copy number per plate), then averaged.

### Statistical methods

All statistics were performed in the R statistical environment (R Core Team 2018). CH_4_ flux and community differences among regions and land types were assessed using a Kruskal-Wallis test, which does not rely on assumptions of distribution and can handle imbalanced sampling designs. Pairwise differences among groups were assessed using Dunn’s test for multiple comparisons. Differences in community structure were assessed with a PERMANOVA test using the ‘adonis’ function in the vegan package in R (Oksanen *et al*. 2015).

Sequence depth per sample ranged from 62,865 to 148,053 sequences per sample, median: 77,653. To account for these differences in sampling depth, the community matrix was rarefied to 62,800 counts per sample ten times and averaged, which did not exclude any samples. The rarefied community matrix was also subsetted for known methanogens and methanotrophs (Supplementary Table 2). We compiled a table of microbial community attributes that represent putative controls on CH_4_ emissions, including abundance, diversity, and composition (Table 1).

**Table 1:** Microbial community attribute measurements used to identify relationships between communities and CH_4_ flux. ASV = Amplicon sequence variant.

| Attribute | Variables |
| --- | --- |
| Abundance | qPCR (of *mcrA* & *pmoA* genes)<br>Relative abundance (in prokaryotic 16S rRNA gene - based community) of methanogens/methanotrophs |
| Composition | Matrix of 16S rRNA gene-based prokaryotic community<br>Principle Components (PCs) 1, 2, 3, 4, 5, & 6 of 16S rRNA gene community matrix |
| Diversity | ASV richness of prokaryotic community<br>Shannon (ASV) diversity of prokaryotic community<br>ASV richness of methanogen/methanotroph community subsets<br>Shannon diversity of methanogen/methanotroph community subsets |

We tested for a relationship between the relative abundance of each taxon and CH_4_ flux. To account for systematic differences in taxon relative abundances due to species interactions, local environmental selection, dispersal history between sites, and other factors unrelated to CH_4_ dynamics, we performed a principal components (PC) correction using the community, environmental, and spatial variables with the ‘prcomp’ function in R (Price *et al*. 2006; Morris *et al*. 2019). For the environmental covariates, we included all soil chemical variables that were shared across samples and that had no missing values and scaled them to unit variance (to account for differences in units of measurement). To account for community structure, we performed principal components analysis (PCA) on the rarefied 16S rRNA gene community matrix following Hellinger transformation and after scaling for unit variance. Spatial coordinates (latitude and longitude) of each sample were assigned a PC score following the same procedure. CH_4_ fluxes, the relative abundance of each taxon in the community matrix, and each community attribute were then adjusted by the principal components for each covariate (community, environment, and geography). This principal components correction removed the correlation between CH_4_ flux and community similarity, environmental similarity, and spatial proximity as well as the correlation between taxon relative abundance and each of the covariates, allowing us to test the unique contribution of each taxon to CH_4_ flux independent of these underlying factors (Price et al. 2006; Morris et al. 2019). We regressed each corrected taxon or community attribute against log_10_-transformed CH_4_ fluxes, and applied a Bonferroni correction (alpha = 0.05) to conservatively address the issue of false positives associated with large numbers of comparisons. Taxa significantly correlated with CH_4_ flux were subsetted from the rarefied community matrix and reduced to a single variable using PCA, and then regressed against log_10_-transformed CH_4_ fluxes. Model fit (R^2^) was assessed after confirming normal distribution of residuals. In several instances one or two high leverage outliers, i.e. “influential outliers” (as defined by Aguinis *et al*. 2013), were removed due to their strong and disproportionate influence on model fit (R^2^).

## RESULTS

### CH_4_ flux and microbial community attributes differ across land types

CH_4_ fluxes were significantly different across land types in both regions (Kruskal Wallis Chi-squared = 33.98, df = 5, *p* < 0.001, Fig. 1A). In both regions, pasture soils emitted CH_4_ at higher rates than primary forest or secondary forest soils (Dunn test for multiple comparisons *p* < 0.001). Combining land types from the two regions, the same pattern emerges, i.e. CH_4_ emissions vary by land type (Chisquared = 25.11, df = 2, *p* < 0.001, Fig. 1B) and pastures emit CH_4_ at significantly higher rates (Dunn test *p* < 0.001) than primary or secondary forests. Of the 25 pasture measurements, only one exhibited CH_4_ uptake (−11 μg CH_4_ m^-2^ d^-1^). In Rondônia, all pasture fluxes were positive, with rates ranging from 30 to 40,000 μg CH_4_ m^-2^ d^-1^ (mean =5,695.3 ± 11,860.5 μg CH_4_ m^-2^ d^-1^). In Pará, pasture emissions were lower, ranging from −11 to 400 μg CH_4_ m^-2^ d^-1^ (mean 93.6 ± 157.9 μg CH_4_ m^-2^d^-1^). CH_4_ fluxes in the Rondônia primary forests ranged from −160 to 550 μg CH_4_ m^-2^ d^-1^ (mean = 22 ± 156.2 μg CH_4_ m^-2^ d^-1^), with four of the fifteen measurements exhibiting uptake, three exhibiting near zero fluxes, and eight emitting CH_4_. Six out of the ten measurements in Pará primary forests exhibited CH_4_ uptake, one had a near zero flux, and three had low levels of emission, ranging from −30 to 8 μg CH_4_ m^-2^ d^-1^ (mean −8.6 +/- 13.2 μg CH_4_ m^-2^ d^-1^). Secondary forests in both regions exhibited CH_4_ uptake on average (Rondônia: mean −17.8 ± 37.7 μg CH_4_ m^-2^ d^-1^, Pará: mean −4.9 ± 34.8 μg CH_4_ m^-2^ d^-1^). Flux values for secondary forest soils in Rondônia ranged from −80 to 30 μg CH_4_ m^-2^ d^-1^, while fluxes from secondary forest soils in Pará ranged from −54 to 61 μg CH_4_ m^-2^ d^-1^.

**Figure 1:**
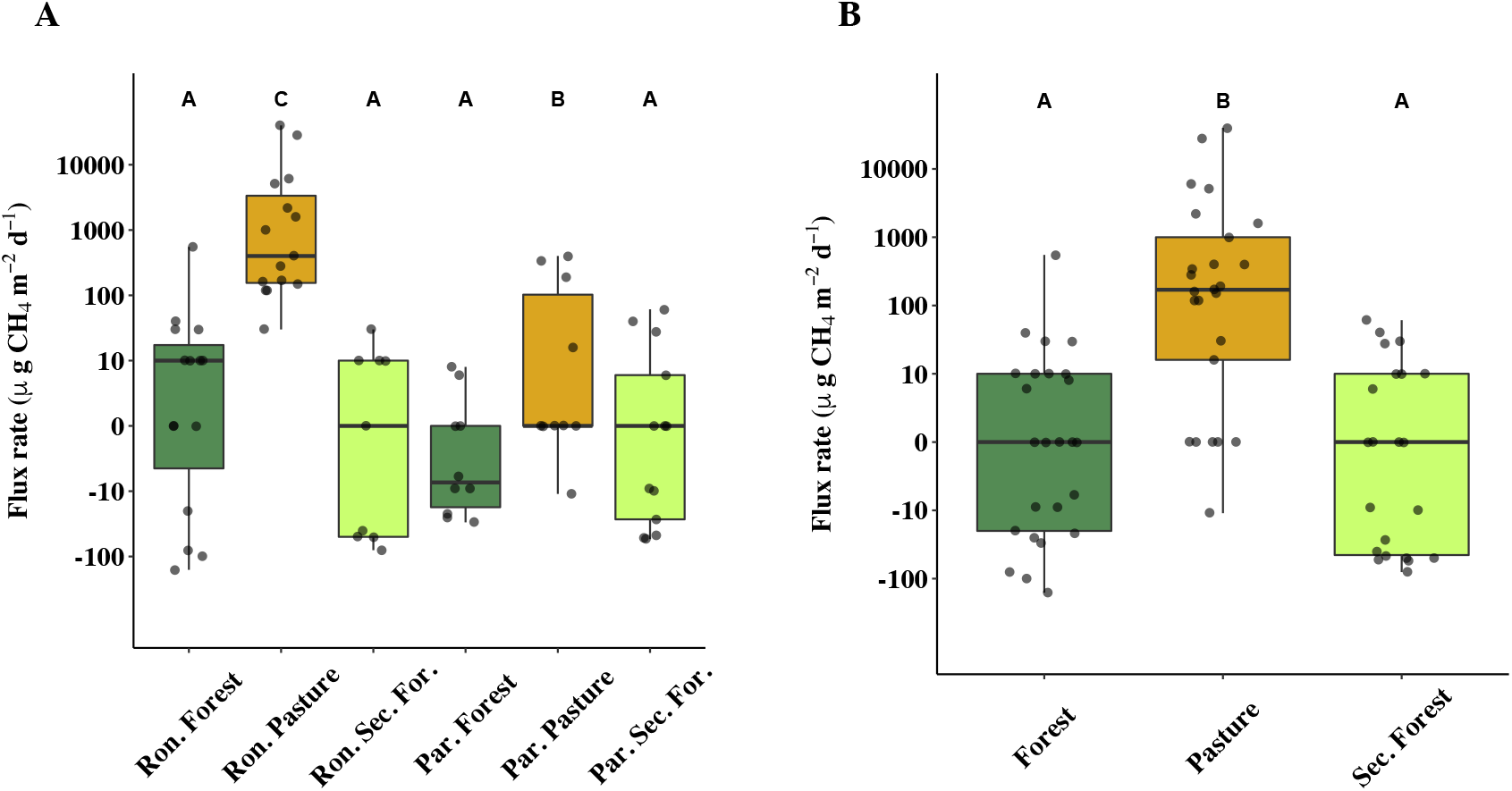
Increased rates of CH_4_ emission in cattle pasture relative to primary forest and secondary forest. Note the log_10_ scale of y-axis values. A) CH_4_ emission rates in forest, cattle pasture, and secondary forest (Sec. For.) across two regions of the Amazon Basin: Rondônia (Ron.) and Pará (Par.). B) CH_4_ fluxes by land type (both regions combined). Pairwise differences between groups (letters A, B, C) were determined using Dunn’s test of multiple comparisons with *p* < 0.05 as significance cutoff.

Our taxonomic survey identified 30,809 prokaryotic (bacterial and archaeal) amplicon sequence variants (ASVs) across the 72 soil samples. Prokaryotic community structure differed by land type (i.e. primary forest, cattle pasture, or secondary forest, PERMANOVA on Bray-Curtis dissimilarities: F_2,68_ = 9.4, R^2^ = 0.18, *p* < 0.001), as well as by region (i.e. Rondônia vs Pará, F_1,68_ = 15.6, R^2^ = 0.15, *p* < 0.001, Supp. Fig. 1). Taxonomic richness (ASV level) also differed by region and land type (Chi-squared = 54.07, *p* < 0.001, Supp. Fig. 2). Across regions, richness values were higher in Rondônia (across all three land types) than Pará (all comparisons Dunn test *p* < 0.001). In Rondônia, cattle pastures and secondary forests had significantly higher richness than primary forests, while in Pará, pastures were the richest, and primary and secondary forests were statistically indistinguishable, but lower than pasture.

**Figure 2:**
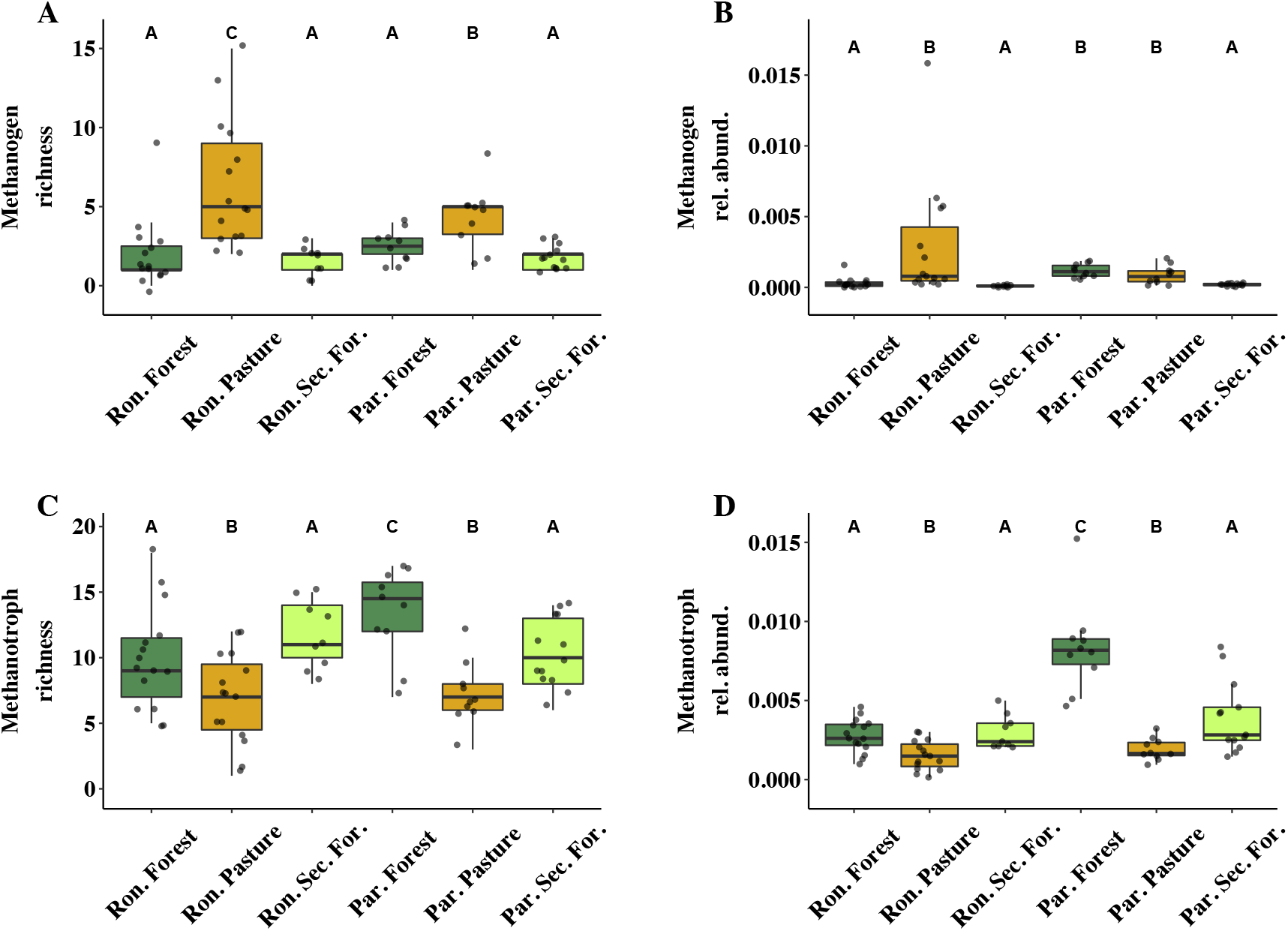
CH_4_-cycling taxa response to land use change in two regions of the Amazon: Rondônia (Ron.) and Pará (Par.). A) The ASV-level taxonomic richness of methanogens by region and land type (inferred from 16S rRNA gene sequences). B) The relative abundance of methanogens in the 16S rRNA gene-inferred prokaryotic community across land use and region. C) The ASV-level taxonomic richness of methanotrophs (inferred from 16S rRNA gene sequences). D) The relative abundance of methanotrophs in the 16S rRNA gene-inferred prokaryotic community. Pairwise differences between groups (letters A, B, C) were determined using Dunn’s test of multiple comparisons with *p* < 0.05 as significance cutoff. Sec. For. = Secondary forest.

Land use change drove numerous alterations to the diversity, abundance, and composition of CH_4_-cycling communities. Methanogen ASV-level richness significantly increased in pastures relative to forest and secondary forest in both regions (Chi-squared = 28.86, df = 2, *p* < 0.001, Fig. 2A). The abundance of methanogens (copies of *mcrA* per g soil) varied by land type and region (Chisquared = 45.62, df = 5, *p* < 0.001), and was higher in pastures relative to primary forests (Rondônia: z = −4.24, *p* < 0.001, Pará: z = −3.91, *p* < 0.001) and secondary forests of Pará relative to pasture (z = −4.37, *p* < 0.001). A similar trend was observed for methanogen relative abundance (in the 16S rRNA gene-derived community) in Rondônia, but in Pará, primary forest and pasture, levels were indistinguishable (*p* > 0.05) and secondary forest abundances were significantly lower than in primary forests or pastures (z = 3.73, *p* < 0.001; z = 2.55 *p* < 0.01, Fig. 2B). Methanogen community composition varied by land type (PERMANOVA on Bray-Curtis dissimilarities: R^2^ = 0.18, *p* < 0.001) and region (R^2^ = 0.06, *p* < 0.001). Most notably, the genera *Methanocella*, *Methanobacterium*, and *Methanosarcina* were almost exclusively detected in cattle pastures of both regions. The genus *Methanomassiliicoccus* varied by land type and region (Chi-squared = 29.18, df = 5, *p* < 0.001), driven primarily by high abundances in the primary forest sites of Pará (*p* < 0.001 for all comparisons).

Methanotroph ASV-level richness also varied by land use (Chi-squared = 18.03, df = 2, *p* < 0.001), decreasing from primary forest to pasture in both Rondônia and Pará (z = 2.01, *p* = 0.02; z = 3.49, *p* < 0.001, respectively). Secondary forest values of methanotroph richness in Rondônia recovered to a level that was statistically indistinguishable from forest (*p* > 0.05), while in Pará levels were higher than in pastures (z = 2.78, *p* < 0.01), but still lower than primary forests (z = −1.76, *p* = 0.04, Fig. 2C). Methanotroph relative abundance was significantly lower in pasture than forest in both Rondônia and Pará (z = 2.45, *p* < 0.01; z = 4.64, *p* < 0.001, respectively). In Rondônia, secondary forest methanotroph relative abundance levels were indistinguishable from primary forest, whereas in Pará levels were significantly lower than primary forest (z = −2.56, *p* < 0.01), but higher than pasture (z = 2.37, *p* < 0.01, Fig. 2D). Methanotroph abundance estimates derived from qPCR of the *pmoA* gene showed a similar, but less pronounced trend across regions and land types (Chi-squared = 10.87, df = 5, *p* = 0.05). Methanotroph composition varied by land use (R^2^ = 0.16, *p* < 0.001), and by region (R^2^ = 0.06, *p* < 0.001). Most notably, the relative abundance of the genera *Methylocella* (Alphaproteobacteria) and *Methylogaea* (Gammaproteobacteria) were significantly lower in pastures relative to forest in both regions (*Methylocella*: Rondônia: z = 3.6, *p* < 0.001; Para: z = 2.13, *p* < 0.05; *Methylogaea*: Rondônia: z = 3.51, *p* < 0.001; Pará: z = 3.86, *p* < 0.001), and increased in secondary forests (*Methylocella*: Rondônia: z = −3.85, *p* < 0.001; Pará: z = −1.65, *p* < 0.05; *Methylogaea*: Rondônia: z = −2.67, *p* < 0.01; Pará: z = −1.91, *p* < 0.05). Lastly, the proportion of methanotrophs in the CH_4_-cycling community (i.e. methanotroph relative abundance divided by the combined relative abundances of methanotrophs and methanogens) was lower in pastures in both regions, but this was only significant in Rondônia (z = 4.71, *p* < 0.001), and secondary forest levels were higher than pasture levels in both regions (Rondônia: z = −5.51, *p* < 0.001; Pará z = −3.22, *p* < 0.001).

### Microbial abundance and diversity are associated with CH_4_ flux

We first asked whether measurements of abundance or diversity (of CH_4_-cycling taxa or the community as whole) could explain variance in CH_4_ flux after accounting for sample covariance structure. Among the best predicting attributes were the ASV-level richness (R^2^ = 0.42, *p* < 0.001, Fig. 3A) and relative abundance (R^2^ = 0.42, *p* < 0.001, Fig. 3B) of methanogens. These were both positive relationships, whereby sites with more abundant and/or diverse populations of methanogens tended to emit CH_4_ at higher rates. The proportion of methanotrophs in the CH_4_-cycling community was negatively associated with CH_4_ flux (R^2^ = 0.36, *p* < 0.001, Supp. Fig. 3); however, this relationship was no longer significant after accounting for covariance structure (*p* = 0.07). No other methanotroph community attributes were related to CH_4_ flux, despite exhibiting strong changes across sites. The only environmental variables significantly associated with CH_4_ flux were pH (R^2^ = 0.08, *p* < 0.05), Zn (R^2^ = 0.21, *p* < 0.001), and Mn (R^2^ = 0.20, *p* < 0.001), all exhibiting positive relationships.

**Figure 3:**
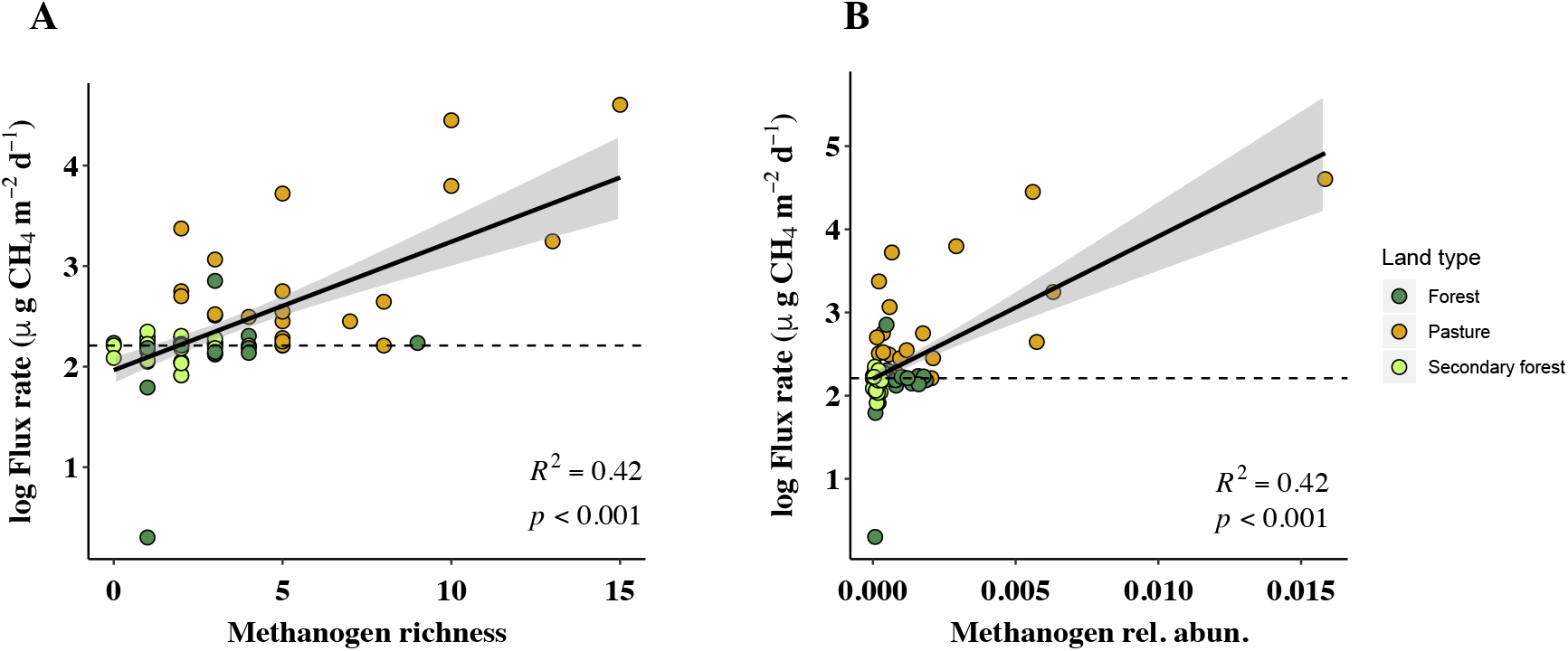
Changes to the A) diversity and B) relative abundance of methanogen taxa are significantly associated with CH_4_ flux across land types and regions, even after accounting for sample covariate structure. R^2^ values represent the proportion of CH_4_ flux variance explained by methanogen attribute, using a linear model on log_10_ transformed CH_4_ flux data. Y-axis is log_10_ transformed with the minimum value added (+162). Dashed line indicates 0 μg CH_4_ m^-2^ d^-1^ flux rate.

### Taxa associated with CH_4_ flux in each land type

We next sought to identify taxa associated with CH_4_ fluxes independent of environmental, spatial, and community covariance structure. We performed our analysis on two datasets: 1) subsets by land type (i.e. primary forest, pasture, and secondary forest) to ask if emissions are controlled by different community members across land types, and 2) across all samples combined. In forest sites we identified 41 (Supp. Table 3) ASVs that together explained 55% of the forest CH_4_ flux variance (*p* < 0.001). None of the taxa are canonically associated with CH_4_ cycling. These taxa included one member of the Thaumarchaeota (Nitrosphaera), and members of eight bacterial phyla, including Acidobacteria, Actinobacteria, Chloroflexi, Firmicutes, Gemmatimonadetes, Planctomycetes, Proteobacteria (divisions Alpha, Beta, Delta, and Gamma), and Verrucomicrobia.

526 taxa across 25 phyla (Supp. Table 4) were associated with pasture CH_4_ fluxes. Only 9 of these taxa are known to directly cycle CH_4_, including 6 methanogens belonging to the genera *Methanocella*, *Methanobacterium*, and *Methanomassiliicoccus*, and 3 Gammproteobacteria methanotrophs belonging to the genera *Methylobacter*, *Methylocaldum*, and *Methylococcus*. Two members of the Crenarchaeota (genus *Thermofilum*) were also among the taxa selected. Collectively the 526 taxa explained 87% of pasture emission variance (regression of subsetted PC1, *p* < 0.001), following removal of one high leverage outlier. There was no overlap at the ASV level between the taxa identified for the primary forest and the pasture sites.

For the secondary forest sites, no taxa passed the *p* value cutoff from our Bonferroni correction (*p* < 3.95 x 10^-6^). We relaxed this threshold to *p* < 0.001 and identified six taxa (Supp. Table 5), including a member of *Acidobacteria* group 13, and members of the genera *Gaiella* (Actinobacteria), *Actinallomurus* (Actinobacteria), *Rhodoplanes* (Alphaproteobacteria), *Nitrospirillum* (Alphaproteobacteria), and *Desulfacinum* (Deltaproteobacteria). None of these taxa were associated with primary forest fluxes and only one (*Gaiella*) was associated with pasture fluxes. Collectively, these taxa when reduced to a single variable explain 38% of the CH_4_ flux variance in secondary forests (*p* = 0.001).

### Taxa associated with CH_4_ flux across land types

Lastly, we performed the above-detailed procedure across all three land types in both regions and identified 654 taxa associated with CH_4_ flux (Supp. Table 5). We subsetted all significant taxa from the community matrix, ordinated them, and regressed their PC1 against CH_4_ flux, and the resulting model explained 50.0% (*p* < 0.001) of the CH_4_ flux variance after removal of one high leverage sample (Fig. 4). Many taxa identified were found in the pasture subset, indicating that pasture samples have a large influence over which taxa are chosen. Eleven methanogen taxa were identified, including members of the genera *Methanocella*, *Methanobacterium*, *Methanosarcina*, and *Methanomassiliicoccus*, comprising 1.7% of identified taxa. Four methanotroph taxa were identified, including members of the genera *Methylocystis* (Alphaproteobacteria), *Methylobacter, Methylocaldum*, and *Methylococcus* (all in the Gammaproteobacteria), together comprising 0.6% of taxa identified. However, the majority of taxa identified are not known to directly cycle CH_4_. Six (0.9%) of the taxa identified are members of the acetogenic genera *Acetonema* (Firmicutes), *Thermacetogenium* (Firmicutes), *Clostridium* (Firmicutes), *Sporomusa* (Firmicutes), and five members of the acetic acid bacteria family Acetobacteraceae. We also identified a member of the anaerobic iron-reducing genus *Geothrix* (Acidobacteria, family Holophagaceae). Lastly, six (0.9%) of the identified taxa play roles nitrogen cycling, including members of the diazotroph genus *Nitrospirillum* (Alphaproteobacteria), a member of the genus of denitrifying bacteria *Denitratisoma* (Betaproteobacteria), ammonia oxidizers from the genera *Nitrosospira* (Betaproteobacteria) and *Nitrosococcus* (Gammaproteobacteria), and members of the nitrite-oxidizing genera *Nitrospira* (Nitrospirae) and *Nitrolancea* (Chloroflexi).

**Figure 4:**
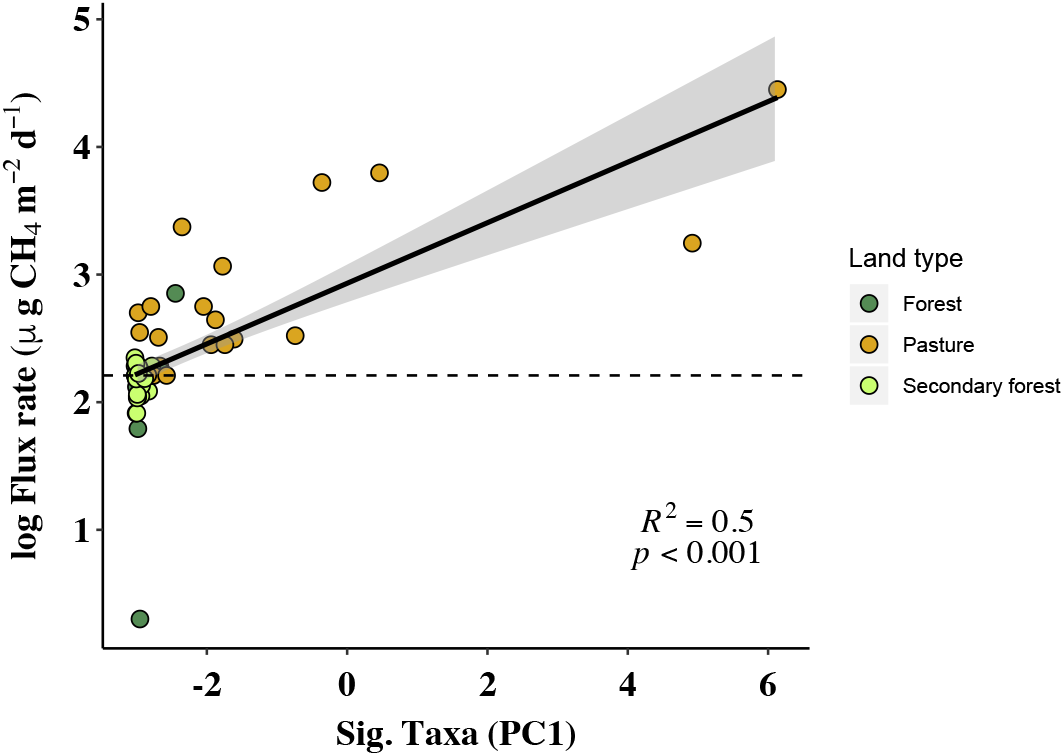
CH_4_ flux is related to a subset of highly associated taxa. Position of points on the X-axis represents the Principle Component 1 (PC1) score representing the 654 taxa that were identified to be highly associated with CH_4_ fluxes, after accounting for sample covariance. R^2^ values represent the proportion of CH_4_ flux variance explained by methanogen attribute, using a linear model on log_10_ transformed CH_4_ flux data. Y-axis is log_10_ transformed with the minimum value added (+162). Dashed line indicates 0 μg CH_4_ m^-2^ d^-1^ flux rate.

## DISCUSSION

Microbial communities drive biogeochemical cycles, including the CH_4_ cycle, but understanding how environmental change influences this relationship remains a crucial challenge. Our results suggest that alterations to microbial community structure resulting from land use change are driving alterations to soil CH_4_-cycling dynamics in Amazon rainforest soils, and thus play a role in the switch from CH_4_ sink to source, as well as the recovery following land abandonment and secondary forest regeneration.

The identity of community members can be an important determinant of ecosystem function (Wardle *et al*. 2011; Díaz *et al*. 2016; Bannar-Martin *et al*. 2018), particularly when species differ in physiological traits such as resource use, allocation, and acquisition (Malik *et al*. 2020). Functional differences among communities can arise when the arrival or persistence of optimal taxa or traits is restricted spatially or temporally (e.g. through dispersal limitation, environmental filtering, or differences in community assembly history). Our results provide compelling evidence for compositional control on the CH_4_ cycle. For example, we identified several methanogens and methanotrophs that were highly associated with CH_4_ flux, suggesting that these taxa disproportionately influence CH_4_ cycling. This included methanogens in the *Methanobacteria*, *Methanocella*, and *Methanosarcina*; all of which increased in relative abundance in pastures relative to forested sites. The methanotrophs identified by our approach also exhibited considerable variation across land types and could influence the flux of CH_4_. For instance, pastures showed increased relative abundance of the genus *Methylocaldum*, and decreased relative abundance of the genus *Methylococcus*. These taxa are known to differ from other methanotrophs in traits related to competitive ability and disturbance tolerance (Ho *et al*. 2013; Knief 2015). Although it is not possible to assess the traits of these organisms from our taxonomic survey, our results suggest that a better understanding of the characteristics of these taxa could improve predictions of CH_4_ cycling.

The majority of the taxa we identified as associated with CH_4_ flux are not known to directly cycle CH_4_, highlighting the importance of considering CH_4_-cycling organisms in a broader community context. Methanogens utilize and compete for metabolic byproducts, including H_2_ and acetate (Westermann *et al*. 1989; Hedderich & Whitman 2013). Six of the taxa associated with CH_4_ flux belong to known acetogenic genera (Müller & Frerichs 2013), which is consistent with suggestions that syntrophic interactions could regulate the CH_4_ cycle (Conrad 1996). We also identified several taxa that could impact the thermodynamic favorability of methanogenesis, or the nutritional demands of methanotrophs. For instance, the production of NO_3^-^_, NO_2^-^_, or the reduction of Fe (III) to Fe (II) are known to limit methanogenesis (Cord-Ruwisch *et al*. 1988; Chen & Lin 1993; Klüber & Conrad 1998; Reiche *et al*. 2008). We identified members of the ammonia oxidizing genera *Nitrosospira* (Betaproteobacteria) and *Nitrosococcus* (Gammaproteobacteria), members of the nitrite-oxidizing genera *Nitrospira* (Nitrospirae) and *Nitrolancea*(Chloroflexi), and the iron-reducing and manganese-reducing genus *Geothrix* as important markers for CH_4_ flux. N-cycling activity could impact the activity of methanotrophs by providing nutrients required for growth (Bodelier *et al*. 2000; Bodelier & Steenbergh 2014). We also identified denitrifier and diazotroph taxa as important predictors of CH_4_ flux, underscoring the interdependence of the C and N cycles. Taken together, these findings suggest that CH_4_ flux in this system could depend on changes to the thermodynamic favorability of methanogenesis as influenced by the activity of taxa involved in redox processes and/or changes to nutrient availability from other community members.

Beyond compositional controls, our results suggest that changes in methanogen abundance and diversity could also be driving increased CH_4_ fluxes in cattle pasture. Methanogen abundance and diversity levels were higher in cattle pasture, which is consistent with another study using metagenomics (Meyer *et al*.2017). This suggests that the soil environment of pastures could be favorable for methanogenesis, perhaps due to an additional supply of labile carbon from grass root exudates and/or decreased O_2_ concentrations throughout the soil column due to compaction (Fernandes *et al*. 2002). Methanogenesis has been positively associated with methanogen abundance and diversity in Congo Basin wetland soils (Meyer *et al*. 2019) as well as anaerobic digesters (Sierocinski *et al*. 2018), suggesting that abundance- and diversity-controls may be common in the CH_4_ cycle.

Our study supports past findings that land use change impacts methanotrophs (Knief *et al*. 2005; Singh *et al*. 2007; Meyer *et al*. 2017), but how these community changes influence CH_4_ flux is less clear. We observed a negative correlation between the proportion of methanotrophs and CH_4_ flux. However, after controlling for environmental variation, this relationship was no longer significant, suggesting that the influence of methanotrophs on CH_4_ flux depends on environmental conditions. Importantly we cannot ascertain whether the methanotrophy process is altered by land use change, as our measurements of CH_4_ flux are the net result of both methanogenesis and methanotrophy. One possibility is that the changes to methanotroph communities that we observed do predict CH_4_ oxidation rates, but that methanotrophy largely does not control CH_4_ fluxes relative to methanogenesis or other processes. Methanotrophy rates have been shown to only predict CH_4_ fluxes when soils are dry and CH_4_ fluxes are negative (Von Fischer & Hedin 2007).

Our study uncovered several relationships between CH_4_ fluxes and soil chemical variables, but the majority of soil chemical variables were not predictive. We saw positive relationships between CH_4_ flux and total soil Zn and Mn levels. Zn plays an important role in the activation of methyl-coenzyme M, a key intermediate for CH_4_ production by all methanogens (Sauer & Thauer 2000). Increased Zn levels have also been shown to stimulate CH_4_ production in tropical alluvial soils under rice production (Mishra *et al*. 1999). Mn has been shown to stimulate methanogenesis in an anaerobic digester system by acting as an electron donor (Qiao *et al*. 2015), and this has also been shown for Zn (Belay & Daniels 1990). We found a weak positive relationship between pH and CH_4_ flux, which is consistent with several other studies (Ye *et al*. 2012; Wagner *et al*. 2017). The general lack of correspondence between the soil chemistry and CH_4_ flux could result from assessing soil chemistry at too coarse of a scale. Microsite conditions are important for anaerobic processes such as methanogenesis, and it has been suggested that better quantifying soil chemistry at microscales could improve our ability to predict CH_4_ emissions (Von Fischer & Hedin 2007). Future work could take a more refined approach by concurrently measuring chemistry and CH_4_ production at smaller scales.

Our CH_4_ flux results provide a sobering look into a potential feedback between climate and land use change. In both regions that we studied, cattle pastures were sources of CH_4_ to the atmosphere. Steudler et al. (1996) and Fernandes et al. (2002) were the first to document the CH_4_ sink-to-source transition of Rondônia soils following forest-to-pasture conversion. These studies reported pasture emissions as high as 0.52 mg CH_4_-C m^-2^ h^-1^ (12,480 μg CH_4_ m^-2^ d^-1^, converted to the units of this study) and 614 mg CH_4_-C m^-2^ yr^-1^ (1682.2 μg CH_4_ m^-2^ d^-1^), respectively. The maximum rate we observed was 40,000 μg CH_4_ m^-2^ d^-1^, and our average CH_4_ flux rate across Rondônia pastures was 5,695.3 μg CH_4_ m^-2^ d^-1^. Our pasture emission estimates are therefore substantially higher than past estimates in the same region. The highest rates of CH_4_ consumption in forest soils from Steudler et al. (1996) were during the dry season, where the maximum uptake rate was 0.061 mg CH_4_-C m^-2^ h^-1^ (1464 μg CH_4_ m^-2^ d^-1^), with rates two-fold lower during the wet season. Our highest rate of consumption was roughly an order of magnitude lower (160 μg CH_4_ m^-2^ d^-1^) than Steudler et al. (1996). Importantly, we sampled during the wet season, when uptake rates would be expected to be lower. Nevertheless, the differences between our uptake rates and Steudler et al. (1996) could represent spatial or temporal variability or the indirect effects of habitat fragmentation due to on-going deforestation activities in the region. Taken together, the immense variability of our CH_4_ flux data and the differences between our study and other work highlight the importance of continuing efforts to study the spatio-temporal dynamics of CH_4_-cycling in the Amazon Basin.

In both Rondônia and Pará, we see a recovery of CH_4_ uptake rates in secondary forest, and on average secondary forest soils consume CH_4_ at rates higher than the primary forests we surveyed. This suggests that forest regeneration could return ecosystems to CH_4_ sinks. Our microbial analyses indicate that secondary forest microbial communities begin to resemble primary forest in the composition and diversity of both CH_4_-cycling organisms as well as the broader community. Therefore, pasture abandonment and reforestation could be a viable strategy for climate mitigation and microorganisms seem to be mediating this response. A final consideration across land types is the role that trees may play in the exchange of soil gases produced at depth. Tree-mediated CH_4_ emissions have been reported to comprise a substantial portion of the Amazon CH_4_ budget, particularly in seasonally inundated zones (Pangala *et al*. 2017). Thus, an untested possibility is that the removal of trees could redirect CH_4_ fluxes through the soil and that secondary forest generation may redirect these fluxes through tree tissue.

Ongoing deforestation and forest-to-pasture conversion in the Amazon Basin is resulting in a switch from ecosystems that are net CH_4_ sinks to those that are net CH_4_ sources. Understanding the mechanism for this change is important not only for our fundamental understanding of global biogeochemical cycles but also for how we manage these ecosystems and model future climate impacts of land use change. With the threat of land use change increasing across the Amazon Basin (Barlow *et al*. 2019; Carvalho *et al*. 2019) it is necessary to improve our understanding of the relationship between community change and ecosystem function. We have shown not only that microbial composition is crucial for understanding CH_4_ dynamics, but also that microorganisms provide explanatory power that cannot be captured by easily measured environmental variables.

## Supporting information

Figure and Table legends for Sup. Mat.

## ACKNOWLEDGEMENTS

We thank the owners of Fazenda Nova Vida for access to their land during sampling. Funding for this research was provided by the National Science Foundation – Dimensions of Biodiversity (DEB 14422214), NSF-PEER (Award 589), NSF-FAPESP (2014/50320-4), and CNPq-311008/2016-0. We thank W. Piccini and A. Pedrinho for assistance in the field.

