## Supplementary material for "Belowground changes to community structure alter methane-cycling dynamics in Amazonia": Figure and Table legends for Sup. Mat.

**Supplementary Figure 1**: 16S rRNA -inferred microbial community composition differs by region and land type. R^2^ and *p* values are the result of a PERMANOVA test on Bray-Curtis community dissimilarities.

**Supplementary Figure 2**: Per sample amplicon sequence variant (ASV) richness varies by land use (primary forest, cattle pasture, and secondary forest) and region of Amazonia (Rondônia (Ron.) and Pará (Par.). Pairwise differences (indicated by A, B, C) assessed by Dunn’s test for multiple comparisons. Each sample was rarefied to 62,800 observations to account for differences in sampling depth.

**Supplementary Figure 3:** The proportion of methanotrophs in the CH_4_-cycling community (i.e. methanotrophs / [methanotrophs + methanogens]) is negatively correlated with CH_4_ fluxes. Note that the significance of this relationship declined to *p* = 0.07 after accounting for covariance structure.

**Supplementary Table 1:** GPS coordinates, Brazilian state, land type, and soil chemical variables for each sample included in the study.

**Supplementary Table 2:** Taxonomic affiliation of methanogen and methanotroph taxa subsetted from the 16S rRNA gene-inferred community matrix.

**Supplementary Table 3:** Taxonomic affiliation of the taxa that were highly associated with CH_4_ fluxes in the primary forest sites after accounting for spatial, environmental, and community covariate structure.

**Supplementary Table 4:** Taxonomic affiliation of the taxa that were highly associated with CH_4_ fluxes in the pasture sites after accounting for spatial, environmental, and community covariate structure.

**Supplementary Table 5:** Taxonomic affiliation of the taxa that were associated with CH_4_ fluxes in the secondary forest sites after accounting for spatial, environmental, and community covariate structure. Note these taxa were only significant after relaxing the *p*-value cutoff to <0.001.

**Supplementary Table 6:** Taxonomic affiliation of the taxa that were highly associated with CH_4_ fluxes across all sites after accounting for spatial, environmental, and community covariate structure.
