## Supplementary figures and images for "Belowground changes to community structure alter methane-cycling dynamics in Amazonia"

### Supplementary Figure 1

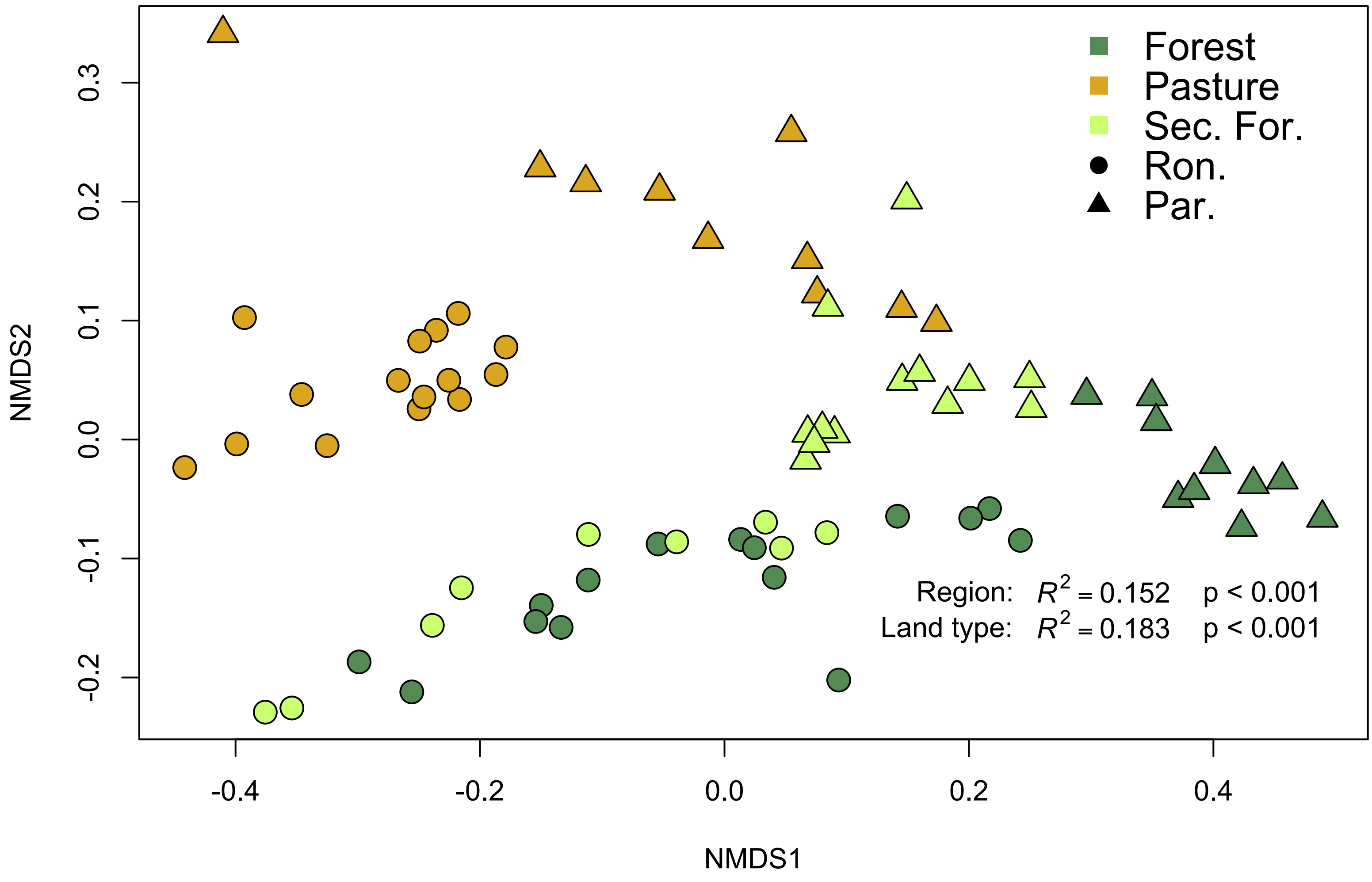

### Supplementary Figure 2

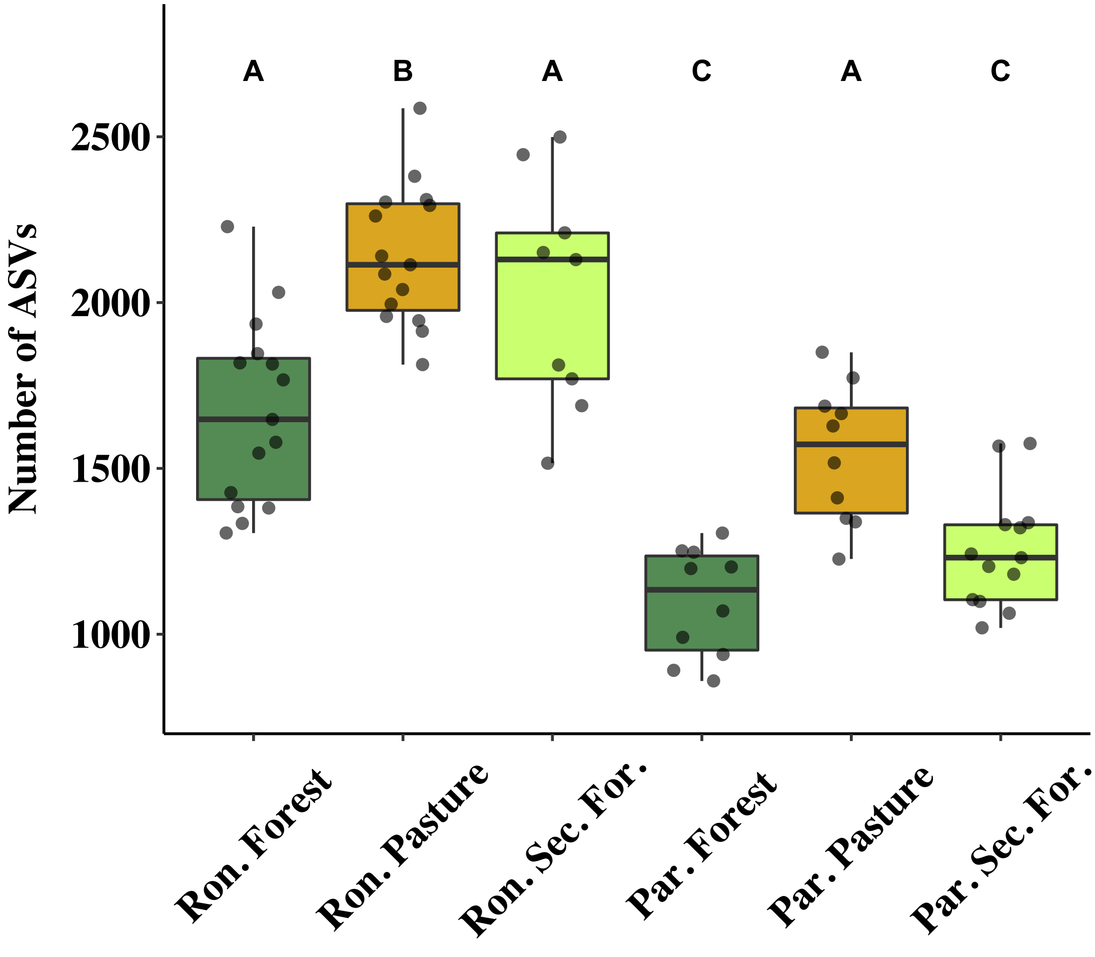

### Supplementary Figure 3

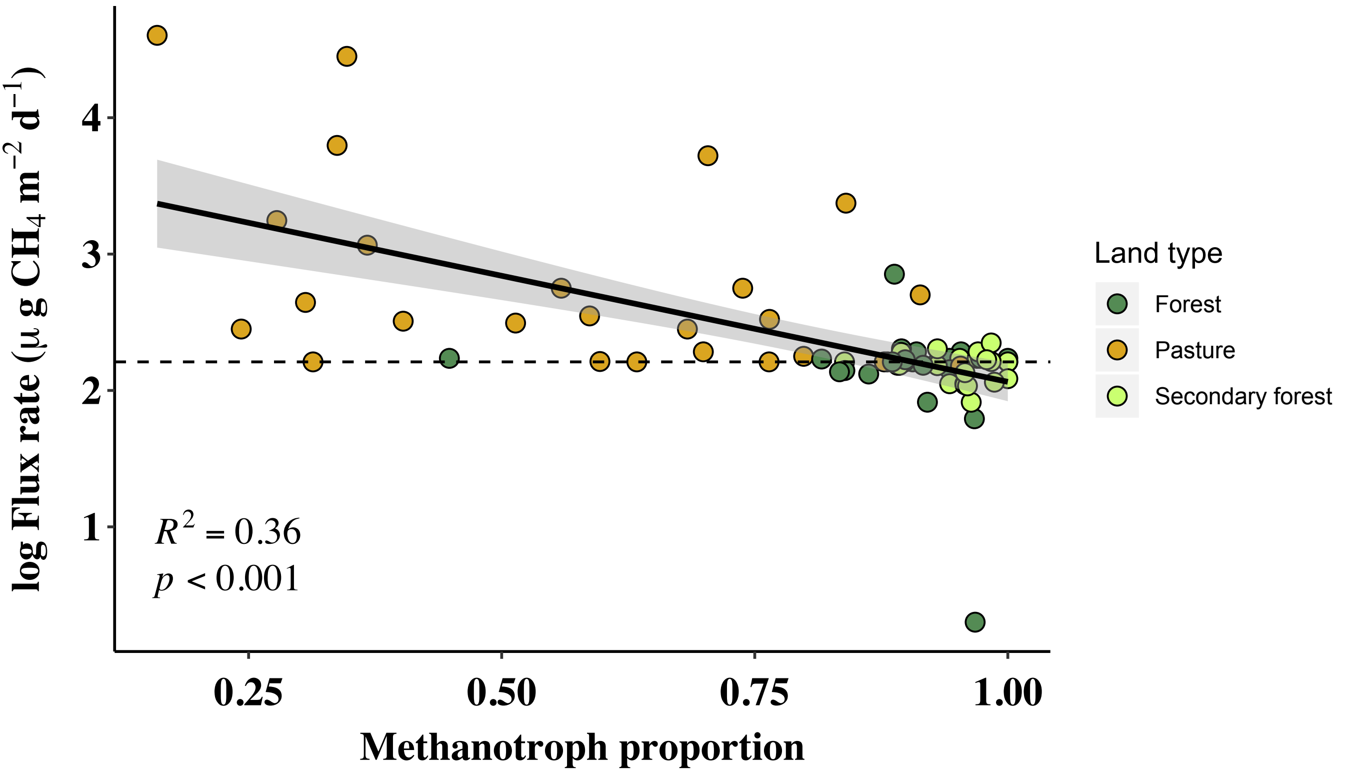
